# Opto-E-Dura: a Soft, Stretchable ECoG Array for Multimodal, Multi-scale Neuroscience

**DOI:** 10.1101/2020.06.10.139493

**Authors:** Aline F. Renz, Jihyun Lee, Klas Tybrandt, Maciej Brzezinski, Dayra A. Lorenzo, Mouna Cerra Cheraka, Jaehong Lee, Fritjof Helmchen, Janos Vörös, Christopher M. Lewis

**Affiliations:** Institute for Biomedical Engineering, ETH Zurich, 8092 Zurich, Switzerland; Laboratory of Neural Circuit Dynamics, Brain Research Institute, University of Zurich, 8057 Zurich, Switzerland; Neuroscience Center Zurich, University and ETH Zurich, Zurich, Switzerland; Laboratory of Organic Electronics, Department of Science and Technology, Linköping University, 601 74 Norrköping, Sweden

**Author notes:** Equal contribution (Janos Vörös, Christopher M. Lewis). Correspondence to JV and CML.

**Keywords:** stretchable electronics, ECoG, in vivo, multimodal recording, PDMS

## Abstract

Soft, stretchable materials hold great promise for the fabrication of biomedical devices due to their capacity to integrate gracefully with and conform to biological tissues. Conformal devices are of particular interest in the development of brain interfaces where rigid structures can lead to tissue damage and loss of signal quality over the lifetime of the implant. Interfaces to study brain function and dysfunction increasingly require multimodal access in order to facilitate measurement of diverse physiological signals that span the disparate temporal and spatial scales of brain dynamics. Here we present the Opto-e-Dura, a soft, stretchable, 16-channel electrocorticography array that is optically transparent. We demonstrate its compatibility with diverse optical and electrical readouts enabling multimodal studies that bridge spatial and temporal scales. The device is chronically stable for weeks, compatible with wide-field and 2-photon calcium imaging and permits the repeated insertion of penetrating multi-electrode arrays. As the variety of sensors and effectors realizable on soft, stretchable substrates expands, similar devices that provide large-scale, multimodal access to the brain will continue to improve fundamental understanding of brain function.

## Introduction

Over the past decade, rapid advances in the field of conformal electronics have brought soft, stretchable materials to the forefront in the design of next generation biomedical devices. In contrast to traditional rigid devices, they integrate well with biological tissues as a result of low mechanical mismatch and high conformability to curved biological structures^[1–3]^. Recently, soft devices have been developed to interface with the central nervous system. Minev et al. introduced the “e-dura”, a soft, stretchable multi-electrode array to address spinal cord injuries^[4]^. Subsequent work improved fabrication techniques and introduced approaches based on nanomaterials^[5]^, hydrogel^[6]^ or inkjet printing^[7]^, and new designs such as meshes^[8]^ and fibers^[9]^. Tybrandt et al. demonstrated the chronic stability of stretchable electrocorticography (ECoG) arrays for long-term electrical measurements from the cortical surface of rats for up to 3 months^[10]^. These developments illustrate the exciting possibilities that soft and stretchable electronics offer for long-term interfacing with the central nervous system. However, while brain function arises from a rich array of physiological signals with diverse dynamics across spatial and temporal scales, most current experimental approaches are restricted to a single measurement modality and scale^[11]^. Understanding brain function requires devices that enable the combined application of multiple tools with complementary resolution and coverage to link the activity of cellular processes, local networks, and patterns of global brain activity. New devices that enable multimodal and multi-scale data acquisition will lead to a more complete picture of brain function and dysfunction.

Experimentally accessible brain signals each have unique strengths and limitations. In isolation they provide a limited, piece-meal picture of brain function. Signals vary in terms of both spatial and temporal scale as well as in the cellular processes they render visible^[12,13]^. ECoG has rapidly become a core method in neuroscience because of its relatively minimal invasiveness, long-term stability, and temporal resolution. However, the neuronal sources that contribute to the ECoG signal are still ill-defined, because ECoG typically lacks the specificity necessary to investigate cellular processes^[14]^. Understanding the ECoG signal is of critical importance for many clinical and basic neuroscience questions. It requires integration with other methods that permit monitoring of identified cell-types based on anatomy, physiology, and genetics^[15]^. Multi-electrodes arrays (MEAs) can be inserted into the brain enabling increased proximity to sources and direct recording of cellular-scale signals within local networks (single- and multi-unit activity, SUA/MUA), as well as population activity similar to ECoG, but more spatially localized (local field potential, LFP). New methods promise to increase the scale and resolution of MEA recordings^[16]^, however, especially for long-term chronic studies, there is a trade-off between dense local sampling and large-scale sampling of distributed populations due to space constraints^[11]^. Optical imaging approaches, such as wide-field and 2-photon calcium imaging, enable cell type and pathway-specificity, as well as long term stability through targeted expression of indicators^[17]^. However, they typically have low temporal resolution, require a trade-off between field of view, resolution and temporal sampling rate^[18]^, and do not provide an absolute measurement because they are derived from changes in fluorescence.

Various attempts have been made to bridge scales by integrating measurement modalities to provide comprehensive perspectives on brain activity^[19–22]^. However, most existing studies have been performed under anesthesia and the vast majority have been performed in acute preparations, where it is impossible to study behavior or learning. Few alternatives exist to provide stable, long-term access for repeated, multimodal, and multi-scale measurement from populations over behaviorally relevant time scales. Such devices will open new windows on integrated brain activity by facilitating the application of complementary tools^[23–25]^. By combining ECoG measurements with complementary optical and electrical methods, we aim to bridge local and distributed dynamics to aid understanding of brain function.

We introduce the ‘Opto-e-Dura’, an optically transparent ECoG array that is chronically stable and facilitates multimodal, multi-scale experiments in combination with diverse optical and electrical methods. The Opto-e-Dura is constructed with a novel fabrication technique, and we demonstrate combined applications of the ECoG with three essential techniques in systems neuroscience: wide-field calcium imaging, 2-photon calcium imaging, and recording from acutely inserted MEAs. The integration of these diverse techniques enables experiments that combine the unique strengths of each method. These applications demonstrate the utility of this new device to improve understanding of brain function in a versatile fashion in the same animal across behaviorally-relevant time scales.

## Results

### Opto-e-Dura design

We developed a PDMS-based ECoG array with highly porous platinum electrodes to interface chronically with the central nervous system based on earlier work^[4,10]^ (**Figure 1**a). We sought to exploit two additional features of PDMS to produce a device for multimodal, multi-scale neuroscience. First, the optical transparency of PDMS makes it suitable as a window to facilitate optical measurement of brain activity^[25,26]^. Second, the soft, elastic nature of PDMS makes it possible to penetrate the membrane with penetrating MEAs and record cellular and population activity from deep and superficial sources^[27]^. We designed a 16-electrode ECoG to match the channel count of our amplifier headstage and to produce a compact package that would fit comfortably on the skull of a mouse. A compact design was especially crucial to permit adequate clearance for objective lenses and MEAs to approach the Opto-e-Dura after implantation. The electrode layout consists of two concentric circles of 8 electrodes each, covering a circular region of ∼4 mm in diameter, roughly 1/3 of the dorsal cortex in one hemisphere of an adult mouse (**Figure 1**b). The array can be positioned on the dura to permit access to a number of sensory and association areas commonly engaged in sensory discrimination behaviors^[28]^ (**Figure 1**b). The electrodes are ring-shaped with an inner diameter of 200 µm and an outer diameter of 400 µm. To facilitate chronic optical access, we designed the device to replace a portion of the cranium and integrated a PDMS imaging well to retain fluid for use with a water-dipping 2-photon objective lens (**Figure 1**c). The high conformability of the Opto-e-Dura allowed us to flexibly position the electrical connector to increase clearance around the ECoG while still connecting the necessary recording electronics. The final device provided good optical access to areas of interest (**Figure 1**d), was chronically stable for up to a month, and enabled multiple imaging and electrophysiological recording sessions in awake and anesthetized animals.

**Figure 1.**
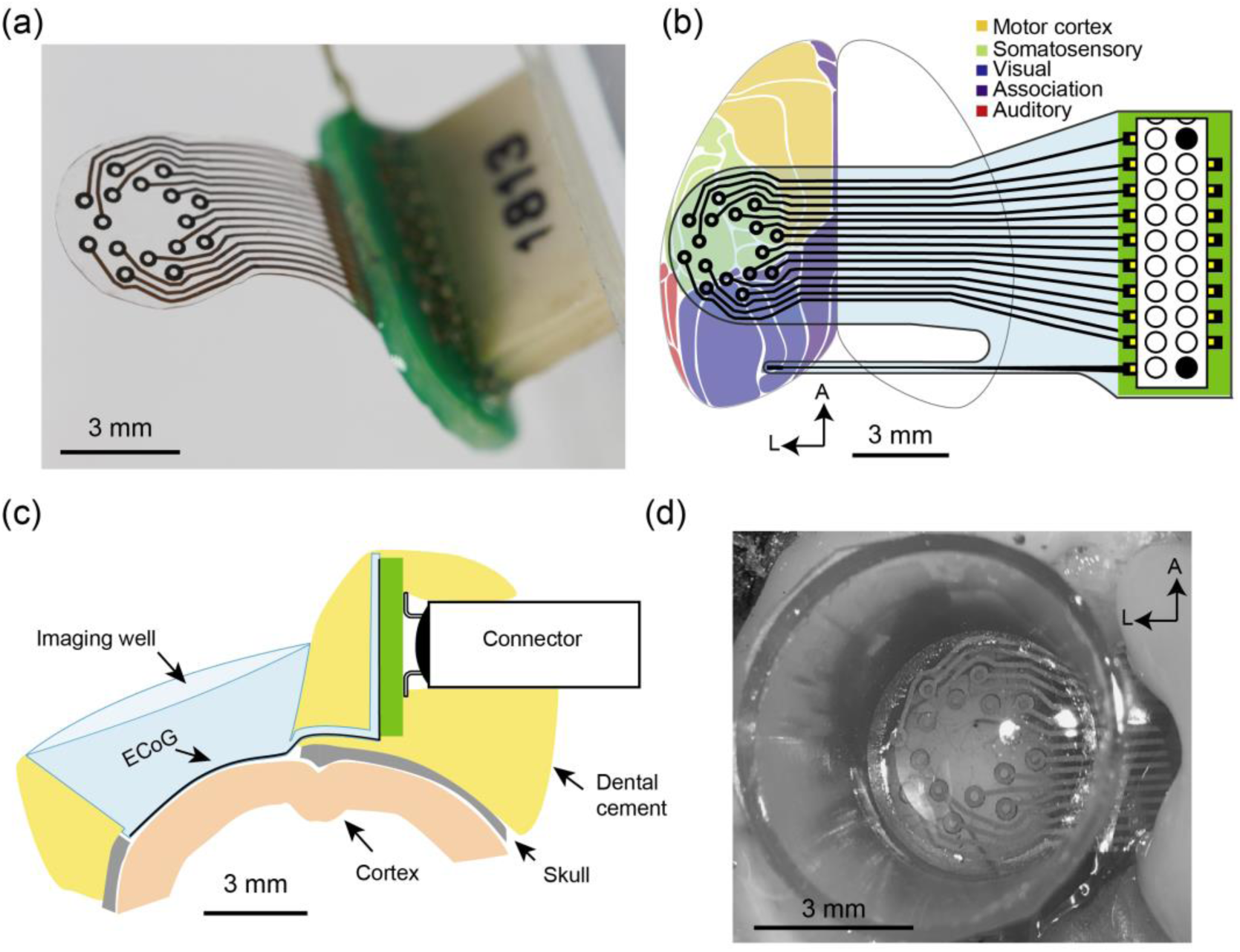
The Opto-e-Dura a) Photograph of the ECoG array before attachment of the imaging well b) Schematic of the device showing its size and coverage over regions of interest on the dorsal cortex of the mouse brain, c) Schematic cross section of the device implanted on the mouse illustrating imaging well and placement of connector to facilitate access of microscope objective, d) Photograph of a device implanted over the dorsal cortex of a mouse.

### Opto-e-Dura fabrication

The ECoG was fabricated using PDMS as a substrate and by combining gold nanowire (Au NW) tracks with platinum (Pt) particle electrodes in a novel push-through approach (**Figure 2**a). The process consists of three main steps: photolithography patterning of filter membranes to generate masks, filtration of conductive components through these masks, and embedding of the conductive patterns in PDMS. First, we performed photolithography on polyvinylidene fluoride (PVDF) filter membranes^[29]^ to generate two masks: one for the Au NW tracks, and one for the Pt particle electrodes (**Figure 2**a). Second, the masks were sequentially placed on a filtration setup to filter a solution consisting of distilled water with either Au NWs or Pt particles through the respective mask (**Figure 2**a). The concentration of the Pt particle solution can be tuned to control the electrode height (Figure S1). By using filtration instead of deposition, we reduce material waste while maintaining high-resolution features^[30]^. Finally, the deposited Au NW tracks are embedded in a semi-cured PDMS layer after careful inspection to insure the absence of defects or short circuits. Crucially, the 2^nd^ PDMS layer must be cured to a pre-defined point for successful electrode fabrication: if it is not cured long enough the Pt particles will diffuse into and be encapsulated by the PDMS, whereas if it is cured for too long it is not possible to achieve electrical contact between the electrodes and tracks. After aligning the membrane containing the Pt particle electrodes with the Au NW tracks on the PDMS, the Pt particle pillars were pushed through the 2^nd^ PDMS layer with a uniform weight (**Figure 2**a) to bring the Pt particle electrodes into electrical contact with the Au NW tracks.

**Figure 2.**
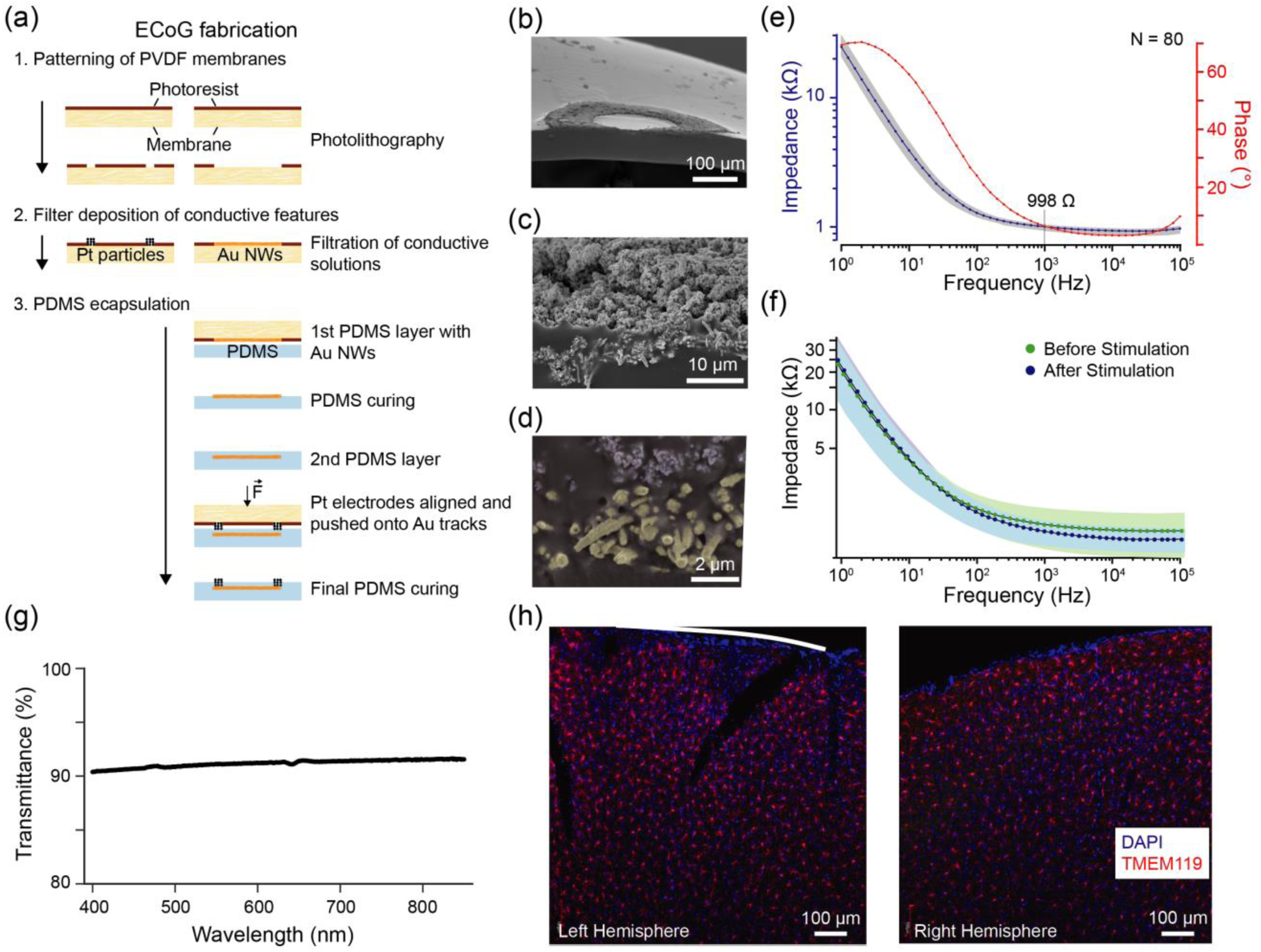
Fabrication and characterization of the Opto-e-Dura. a) PVDF membranes are patterned with photoresist, Pt particles and Au NWs are filtered and sedimented in the negative space, as previously described^[10]^. Au NWs were brought in contact with PDMS before a second layer is added to insulate the tracks. Through the second PDMS layer the Pt particle pillars, acting as electrodes, were brought in contact with the underlying Au NWs by applying force. b-d) SEM images of the side cuts of the whole electrode (400 µm outer and 200 µm inner dimension) (b), the porous Pt particle surface with the underlying Au NWs (c) and the zoomed in Pt particle (blue) to Au NWs (yellow) interface (d). e) The median of the impedance and the bode phase of 80 electrodes (94.3k μm^2^) from 5 of the implanted Opto-e-Dura devices. f) Stimulation stability of 16 electrodes before (green) and after (blue) 10000 biphasic pulses at 200 μA for 200 μs each. g) Spectrophotometry data of a cortex MEA, showing over 90% transmittance in the range suitable for wide field imaging. h) Post-mortem histology of a mouse chronically implanted with an Opto-e-Dura for 6 weeks shows lack of chronic immune response to the implant. DAPI-staining of nuclei (blue) and immune-staining of microglia (red, anti-TMEM119) for a coronal slice through the center of the Opto-e-Dura. No obvious microglia activation is discernible at the location of the array over the left hemisphere (left image) as compared with the control right hemisphere with no implant (right image). White line indicates the position of the Opto-e-Dura.

After testing the functionality of the fully cured device, it was cut to the desired shape and mounted on a PCB with a connector (**Figure 1**a). Scanning electron microscopic (SEM) imaging revealed high porosity of the Pt-particle electrodes (**Figure 2**b) and established contact at the interface between the Au NWs and the Pt particles (**Figure 2**c-d). We fabricated devices of 33μm and 75 μm thickness by adapting the spin coating speed for the first PDMS layer. Here we chose to use the 75 μm devices for further *in vivo* tests to assure both stability during imaging and durability during penetration with linear electrode arrays. However, some applications might require thinner, more conformable devices, were the 33 μm comes in handy and as both thicknesses had similar impedances, there is no drawback in terms of the device’s performance (Figure S2).

### Characterization of electrochemical performance

The impedance spectrum of each device was measured prior to implantation and established the stability and reliability of the fabrication technique. First, cyclic voltammetry (CV) was performed to electrochemically condition the electrodes and establish a stable double layer at the electrode-electrolyte interface. After formation of the double layer, we measured the impedance, which on average was 998 ± 93 Ω at 1 kHz (mean ± SD; N=80 electrodes=). The impedances of the Opto-e-Dura electrodes at high frequencies (>1k Hz), when the resistance is the limiting factor, were consistent with those previously reported for PDMS-based electrodes (**Figure 2**e). However, impedances were consistently lower at lower frequencies than conventional stretchable electrodes(e.g., 24 Ω cm^2^ at 1 Hz compared to 30 Ω cm^2^ in Tybrandt et al.^[10]^). The electrochemical equivalent circuit can be found in Supplementary Figure S3. Contrary to typical impedance properties of metal electrodes, the conductance per unit area significantly increases with decreasing electrode diameter with the presented approach. To the best of our knowledge, this leads to the best results for current PDMS based electrodes with around 3 Ω cm^2^ for 30-µm ring electrodes (Figure S4)^[10]^. Due to the distinct porosity achieved with the push-through technique as seen in Figure 2d, an increased surface area is achieved which has a greater impact the smaller the electrode becomes. This is especially interesting for future studies, when the electrode shape and diameter are not pre-defined to a donut shape by the dimension of linear electrode arrays.

Two different experiments were conducted to test the stability of the device. The impedance spectrum changed negligibly after 10,000 typical stimulation cycles (**Figure 2**f) as well as in an accelerated aging test (Figure S5). Because the layout was transferred by imprinting the patterned membrane onto the PDMS during fabrication, we tested the Opto-e-Dura transparency to ensure that it was suitable for optical imaging. Spectrophotometry revealed over 90% transparency across the visible and near infrared spectrum (**Figure 2**g). Therefore, approximately 10% of both the excitation and emission light is lost leading to a reduced signal to noise ratio. However, these numbers compare favorably with glass and PDMS cranial windows commonly used for imaging^[24]^. Additionally, unlike some materials, such as polyimide, PDMS has exceptionally low autofluorescence in both the visible and near infrared ranges, making it suitable for both wide-field and 2-photon imaging^[24]^.

To establish the chronic biocompatibility of the implant, we performed post-mortem histology on coronal slices of implanted mice to investigate enhanced microglia activation by the array. Slices prepared 6 weeks after ECoG array implantation were stained with DAPI to visualize cell nuclei and immune-stained with an antibody against TMEM119 to label microglia in order to assess the degree to which implantation induced immune reactivity. No qualitatively difference in microglial activation was observed between the implanted and the non-implanted hemisphere (**Figure 2**h).

### Combined ECoG recording and wide-field calcium imaging *in vivo*

To demonstrate the suitability of the arrays for multimodal data acquisition, we first performed simultaneous electrocorticography and wide-field calcium imaging. Arrays were implanted over the dorsal cortex of triple transgenic mice expressing a genetically encoded calcium indicator, GCaMP6f, in excitatory neurons (SNAP25-Cre; CamKIIa-tTA; TITL-GCaMP6f)^[31]^. The animal was positioned to allow optical access through the Opto-e-Dura and wide-field imaging was performed using a standard fluorescence macroscope setup, while electrical signals were digitized and recorded using a commercial electrophysiology system (**Figure 3**a). The transparency of the Opto-e-Dura enabled imaging of the dorsal cortex within a circular field of view (∼4 mm diameter), while simultaneously recording ECoG signals (**Figure 3**b). As expected, the ECoG exhibited faster temporal dynamics than the simultaneously recorded fluorescence (**Figure 3**c), while imaging provided better spatial resolution and coverage, with fluorescence observable across the full field of view. We assessed the spatial and the temporal correspondence between the signals by correlating the band-limited power envelope of the ECoG signal with the percentage GCaMP6f fluorescence changes (ΔF/F) at each imaging pixel. Due to the higher temporal sampling rate of the ECoG compared to the wide-field signal (24 kHz, as opposed to 20 Hz), we first decomposed the ECoG signal into distinct frequency components (1-100 Hz) at the sampling rate of the wide-field imaging camera. This is permissible because, unlike the raw ECoG signal, the envelope of the power typically has considerably slower temporal dynamics^[32]^. With the band-limited power signal from the ECoG, we investigated the spatial (**Figure 3**d), and the spectral distribution (**Figure 3**e) of the covariance between the ECoG and wide-field calcium signals. We found that the calcium signal showed both positive and negative peaks in correlation with the broadband ECoG signal, with positive peaks spatially localized to the region around the reference electrode (**Figure 3**d). We observed additional structure in the spectral domain, as illustrated in the spectrum of the shared variance (**Figure 3**e). These findings suggest intricate coordination across modalities as well as temporal and spatial scales, which demands further investigation.

**Figure 3.**
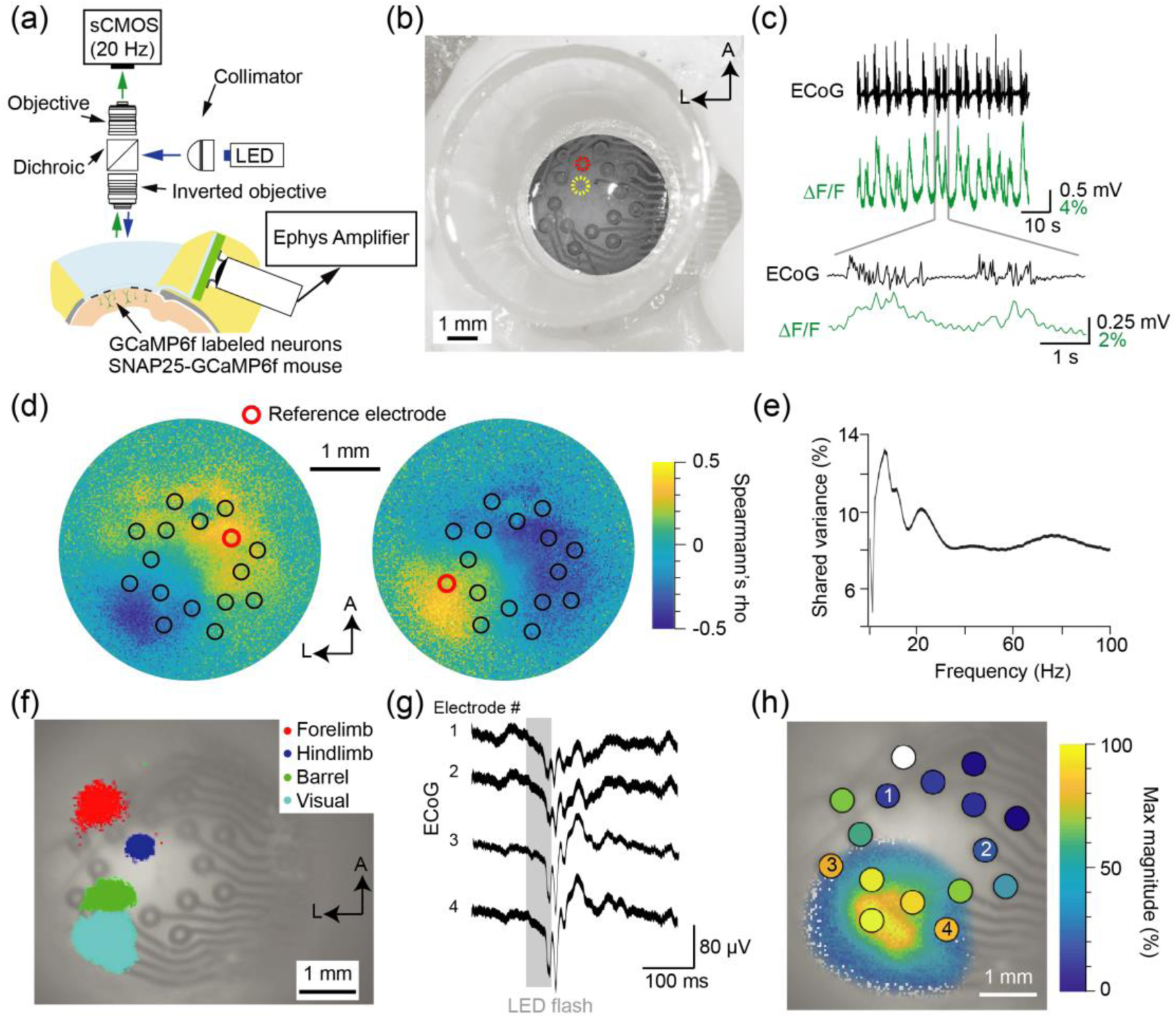
*In vivo* wide-field calcium imaging through Opto-e-Dura. a) Schematic depicting a cross section through the implanted Opto-e-Dura, the wide-field imaging system and the electrophysiology amplifier. b) Photograph of Opto-e-Dura in place over the cortex of a chronically implanted transgenic SNAP25-GCaMP6 mouse. Colored circles indicate locations of example traces in panel c. c) Example of simultaneous ECoG and wide-field fluorescence signals exhibiting correlated dynamics with distinct time scales. ECoG signal is from electrode indicated in red in panel b, and calcium signal is the spatial average of the pixels indicated by the yellow circle in panel b). d) Spatial correlation of the total power in the ECoG signal for two example electrodes (indicated by red circles), with the fluorescence signal at each pixel imaged through the Opto-e-Dura. e) Spectrum of shared variance between the spontaneous calcium signal and the band-limited power of the ECoG signal demonstrating spectral structure of the correspondence. Percent of shared variance was computed using Spearman’s rank correlation. f) Modality specific portions of cortex for four sensory stimuli overlaid on the wide-field image through the Opto-e-Dura. Regions reflect the boundary of peak activity derived from calcium activation maps and are colored to indicate stimulation according to the legend. g) Visually evoked ECoG activity for 4 example electrodes (indicated in panel h) averaged over 20 repetitions (line width indicates standard error of the mean). Stimulus onset and duration indicated by grey shaded box. h) Correspondence of calcium imaging response and simultaneous electrical recordings to the visual stimulus. The response of the electrodes and calcium imaging is normalized and color-coded to illustrate the agreement between the relative amplitudes of the visually evoked potential and calcium responses. The white dot represents a non-functional electrode.

To further establish the correspondence between the signals, we performed functional mapping in anesthetized, head-restrained mice using wide-field imaging while performing ECoG recording. The mapping experiment consisted of blocks of trials stimulating alternate sensory modalities and distinct portions of the body contralateral to the implant: the right whiskers, forelimb, hindlimb and flashes of blue light to the right eye. We were able to localize primary and secondary sensory areas to the expected areas of the dorsal cortex based on the selectivity of the wide-field signal response^[28,33]^ (**Figure 3**f). Sensory responses were also present in the ECoG signal as illustrated by the visually evoked potentials recorded from the Opto-e-Dura (**Figure 3**g). We assessed the similarity in the topography of the visually evoked ECoG and wide-field response by comparing the magnitude of the response across the Opto-e-Dura electrodes and the imaging field (**Figure 3**h). The spatial distribution of the ECoG and wide-field responses exhibited a high degree of correspondence (spatial correlation = 0.83 ± 0.11 mean ± SD, Spearmann rho, N = 3).

### Combined ECoG recordings and 2-photon calcium imaging *in vivo*

While the combination of wide-field calcium imaging and ECoG recording can provide unique information about the contribution of specific cell types to the generation of macroscopic electrical fields, it is ultimately important to achieve single-cell, and even subcellular, resolution. In order to demonstrate the suitability of the Opto-e-Dura for this purpose, we performed 2-photon calcium imaging through the array while simultaneously measuring the ECoG signal. We first modelled the expected loss of excitation and emission light when imaging though the Opto-e-Dura, as a result of opaque elements (electrodes and tracks). As expected, we found that light transmission is reduced for deeper focal planes (**Figure 4**a), resulting in smaller fields of view (Figure S6). In the volume less than 0.4 millimeter below the lower edge of the ECoG, it is possible to image a field of view of approximately 1 mm diameter (Figure S6b). Further losses in signal quality will occur because the ECOG structure destroys the wavefront of the excitation light beam, altering its focal volume and leading to a reduction in resolution, and therefore reduced 2-photon excitation efficiency. These effects can be mitigated by a more complicated excitation path, for example by including a spatial light modulator which negates the destructive effects of the ECoG on the wavefront. It should be noted that the focal-depth-dependent attenuation of excitation and emission light due to ECoG features is independent of the transmission of the PDMS reported earlier. Therefore, both losses will accumulate to reduce the signal to noise ratio. Losses due to the Opto-e-Dura occur in addition to other, common optical losses, such as depth-dependent scattering, an exponential decrease with the tissue-dependent scattering length as parameter.^[34]^

**Figure 4.**
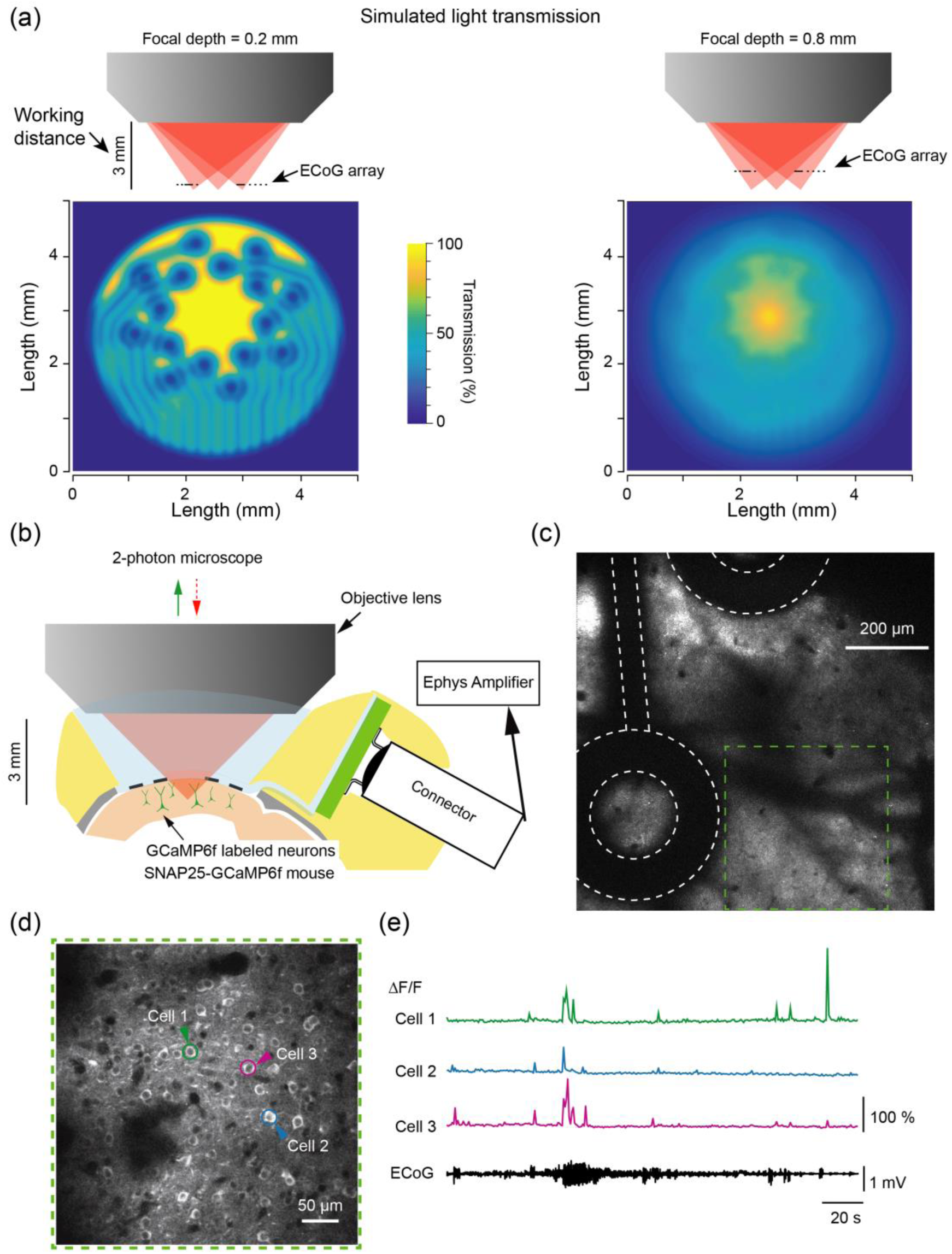
*In vivo* 2-photon calcium imaging through Opto-e-Dura. a) Estimation of signal transmission through the Opto-e-Dura for the 2-photon objective at two focal depths: 0.2 mm (left) and 0.8 mm (right). Top, illustration of excitation beam scanning a plane through the ECoG at two focal depths. Bottom estimated transmission of light at two focal depths. Signal loss is depth dependent. b) Schematic depicting a cross section through the implanted Opto-e-Dura, the 2-photon imaging system, and electrophysiology amplifier. The imaging well is filled with water and placement of the connector and dental cement facilitates access with the objective. c) Large field of view scanned in close proximity to two electrodes indicated by dashed, white outline. Focal depth for the image was at the lower surface of the Opto-e-Dura. Green square region indicates a smaller subfield imaged at a deeper focal depth in d. d) Image of fluorescence from a focal plane 180 µm deeper than to the image presented in panel c, and the bottom of the Opto-e-Dura. Raster scanning was performed in the full field of view in close proximity to the electrode: the left border of the field of view is < 50 µm from the electrode, and 180 µm deeper. Shadows on the left come from the electrodes and the dark spots in the center are blood vessels through the cortex. e) Time series from the cells indicated by colored circles and arrow heads in panel d are presented in with simultaneous ECoG signal. The cells were in close proximity to the electrode: Cell 1 was 200 µm lateral and 180 µm deeper. The time series from the electrode is displayed and correlated events are visible in the traces.

The placement of the ECoG, imaging well, and connector permitted access with the water-dipping 2-photon objective (**Figure 4**b) and it was possible to image in the vicinity of ECoG electrodes and tracks without excessive reflectance or electrical artifact (**Figure 4**c). We did not observe electrical artifacts as a result of scanning the laser across the Opto-e-Dura, unless the laser scanned within 50 µm of a conductive feature of the ECoG, such as an electrode or track. The depth of the imaging plane is also limited by the dimensions of the PDMS well and the working distance of the objective. In our preparation, we were able to image a field of view of size 0.5 mm to a depth of 0.4 mm from the bottom edge of the Opto-e-Dura. It was not possible to image deeper without disturbing the ECoG array, as the objective would approach the sides of the imaging well. More specifically, neurons could be imaged in the vicinity of recording electrodes (within 200 µm lateral from the nearest electrode and 180 µm deeper) while simultaneously measuring ECoG signals (**Figure 4**d). In general, 2-photon calcium imaging through the Opto-e-Dura captured high-resolution single-cell recordings down to several hundred micrometers from the cortical surface, with simultaneous, high temporal resolution, multichannel ECoG signals.

### Combined ECoG and penetrating multi-electrode arrays *in vivo*

In addition to optical and electrical recordings from superficial cortex, an ideal device would enable dense, MEA recordings. Due to the soft and elastic character of PDMS we were able to penetrate the Opto-e-Dura with sharp, MEAs. The combination of ECoG with MEA recording facilitates multi-scale measurements across the large area covered by the Opto-e-Dura and dense recording from the cortex and deep structures. To demonstrate the suitability of the arrays for combined surface and depth recordings, we acutely placed a multi-site linear silicon array through the Opto-e-Dura (**Figure 5**a). Similarly, injection cannulas or micropipettes could be inserted through the ECoG to deliver viruses, dyes, or drugs to the cortex or deeper structures. Here, we coated the linear probe with dye to reconstruct its track through cortex and into hippocampus (**Figure 5**a). We reliably performed multiple insertions (up to 10 times at various locations) of MEAs through the ECoG and recorded activity across multiple cortical and subcortical regions (**Figure 5**b). The maximally achievable depth of insertion is defined by the length and geometry of the array probe, the size of the connector and headstage, and the geometry of the imaging well.

**Figure 5.**
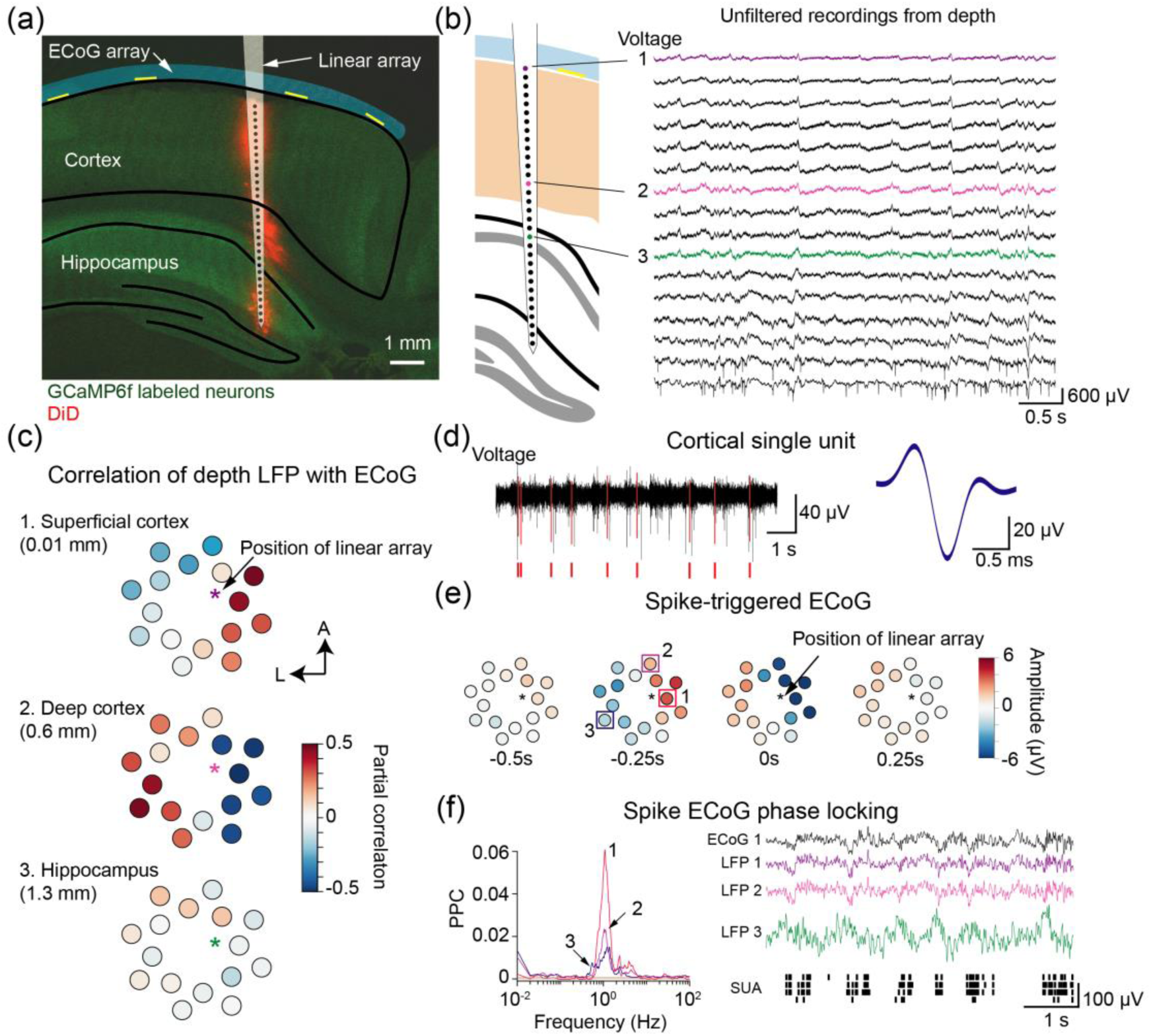
*In vivo* penetration of Opto-e-Dura for simultaneous surface and depth electrophysiology. a) Histological image of a post-mortem slice from a chronically implanted SNAP25 mouse. Green indicates fluorescence from GCaMP6f in neurons expression SNAP25 and red indicates fluorescence from the lipophilic dye DiD. The position of the Opto-e-Dura is depicted and the placement of the acutely placed linear electrode array is illustrated based on DiD labelling. Borders of cortex and hippocampus are overlaid by registration of the histological section with an atlas. b) Schematic of the linear electrode array penetrating the Opto-e-Dura, cortex and hippocampus in close proximity to an ECoG electrode. Example raw voltage traces from acute linear array. c) Spatial maps displayed on the opto-e-dura of the correlation between the ECoG signal and the LFP from the linear array at three depths. Opto-e-Dura electrodes are color coded to indicate the correlation at each electrode with the LFP signal at the indicated depth. As expected, the correlation reverses across cortical sites from superficial to deep and is attenuated for the deepest sites in the hippocampus. d) Single unit activity recorded from cortex on the linear array. Left, high-pass filtered voltage trace from the linear array showing an example single unit indicated by red voltage deflections. Right, the waveform of the single unit indicated on the left. e) Spike-triggered voltage of the ECoG is illustrated spatially across the array for the cortical unit in shown in d. Each ECoG electrode is colored to indicate the average voltage at four time points centered on action potentials from the cortical unit. In the vicinity of the linear array, a spatially localized variation in voltage precedes the spike. f) Phase locking of cortical spikes to the ECoG signal. Left, locking of the single unit shown in d to three example ECoG electrodes estimated by the pair-wise phase consistency (PPC) measure. The cortical unit is strongly locked to slow (∼1 Hz) variations in nearby ECoG electrodes, but the locking breaks down over distance. ECoG electrodes indicated with numbers are displayed on the schematic of the opto-e-dura in panel E. Right, example traces for 10 seconds showing multiple slow network events occurring synchronously across cortex and the hippocampus. ECoG activity is shown along with LFP from the linear array at the locations in panel c, as well as action potentials from 4 isolated cortical single units. Slow events repeat with a periodicity ∼ 1 second, as measured by PPC.

We investigated the correspondence of the ECoG signal to depth recordings for both population activity (LFP) and single cells (SUA). We quantified the spatial and temporal characteristics of the correspondence using correlations, similar to the comparison of the wide-field calcium signal to the ECoG (**Figure 5**c-f). We first quantified how the spatial distribution of ECoG activity was related to the LFP from different depths along the linear array. We performed a partial correlation analysis for each ECoG channel with the LFP from each electrode on the MEA. We found that the correlation showed both positive and negative peaks, with the peak of correlation on the ECoG array occurring in close proximity to the position of the MEA (**Figure 5**c). Further, we found that this relationship varied with depth, diminishing for deeper positions. Finally, we investigated the relationship of the spiking activity of single cortical neurons to the ECoG signal. We isolated single cortical units from the MEA and quantified the spatial distribution of ECoG activity in the time period around action potentials. We found that a wave of surface positivity preceded action potentials, followed by a surface negativity coinciding with the action potential in the vicinity of the MEA (**Figure 5**e). To investigate the spectral characteristics of this synchronization, we estimated the frequency-resolved phase-locking between cortical action potentials and the ECoG signal. We found that cortical action potentials recorded on the MEA were consistently phase-locked to the ECoG signal in a topographical manner, peaking at ECoG sites near the array and falling off with distance The locking had specific spectral structure, peaking around 1 Hz, related to cortical Up-Down state transitions (**Figure 5**f, left). Example traces of simultaneous ECoG, LFP and SUA demonstrate a series of Up-Down state transitions and illustrate additional structure in depth along the MEA (**Figure 5**f, right). Cortical LFP was highly correlated to the proximal ECoG electrode and phase locked to cortical single units, while hippocampal LFP exhibited distinct activity patterns anti-correlated with the profile in the cortex. Combined ECoG and electrode array recordings thus enable multi-scale investigation of local and distributed dynamics, which can be performed repeatedly across days in chronically implanted animals, across behaviorally relevant time-scales.

## Discussion

We designed, fabricated, and validated the Opto-e-Dura, a PDMS-based ECoG array that enables multimodal, multi-scale neuroscience. Using PDMS as a base material, the device is transparent, elastic, biocompatible, and chronically stable for weeks in mice. These features facilitate multimodal experiments that combine ECoG recordings with optical imaging and repeated recordings from penetrating MEAs. The ability to flexibly combine diverse measurement modalities in the same animal enhances the versatility of experimental approaches and facilitates multimodal experiments comprising techniques with complementary strengths. We found rich, highly structured correspondences between the diverse signals we measured at all spatial and temporal scales. Comparisons revealed both spatial and temporal specificity, and pointed future directions for investigating the basis of the ECoG signal. The correspondence between ongoing variations in the time resolved power of the ECoG signal and both the calcium activity and local single unit activity suggests a path towards better understanding this important macroscopic signal in terms of the contributions of specific cell types and even individual cells. Moreover, additional studies could further explore these relationships to provide a better understanding of both the wide-field calcium signal and electrical oscillations in specific frequency ranges by revealing the involvement of specific cellular populations.

Future applications of the opto-e-dura could use the approaches presented here in concert. For example, large-scale functional mapping could be performed using the ECoG and wide-field imaging, and the derived maps could localize regions of interest and their topography. Specific populations could be subsequently selected according to their response properties and targeted for viral injections, drug delivery, or dense recording with 2-photon imaging or MEAs. Such studies enable the simultaneous recording of distributed surface activity, in combination with depth-resolved recordings of targeted populations. Compared with previous attempts at multimodal, multi-scale studies, our approach enables stable, chronic access in awake animals. In chronically implanted animals, experiments can track local and distributed dynamics during the performance of behavioral tasks or the acquisition of novel skills. It is possible to perform repeated imaging or repeated penetrations with MEAs across large portions of the brain without the need for additional invasive surgery or anesthesia. While the presented arrays were robust and chronically stable, enabling electrical and optical access to targeted brain regions for multiple weeks (2-4 weeks), future work will be necessary to improve implantation procedures and the long-term stability of the device, as well as increase the density and decrease the size of electrodes. Additionally, the elasticity of the device can be further leveraged to enable penetration with stimulating electrodes or micro-syringes for targeted injection or drug application. The transparency of the Opto-e-Dura also make it an appealing interface for use with optogenetic activation or silencing studies where light can be delivered to local portions of the cortex, or perturbation experiments where light is raster-scanned across the array to determine regions engaged in a particular task. Overall the Opto-e-Dura demonstrates the evolving potential of stretchable electronics for next-generation brain-machine interfaces.

### Experimental Section

#### ECoG fabrication

Two PVDF membranes were patterned with an established photolithography process (Tybrandt et al. 2016) (microresist technology ma N-490, 4acc, 60s, 3.4 krpm for track and 3 krpm for electrode layer), one for the track layer and one for the electrodes. Subsequently, recently published Au NWs^[10]^ were filtered onto the pre-wetted track layer membrane. For the electrode layer, a 1 bead thick layer of PMMA beads (0.71 µm, Bangs Laboratories Inc.) was filtered prior to the Pt particles (Goodfellow, 3.5 µm) to lower adhesion to the membrane and increase the porosity as they are dissolved after PDMS curing. The bottom PDMS layer of roughly 60 µm thickness (Sylgard184 DowCorning) was spincoated (1 krpm, 30 s) on a silanized borosilicate 2-inch wafer and semi-cured for 3.5 minutes on a hotplate at 75°C before transferring the Au NW layer. After curing an additional 10 minutes at 75°C, the second PDMS layer with a thickness of around 15 µm (5.5 krpm, 60 s) was spin coated on top. This layer was pre-cured for 3 minutes at 75°C to permit penetration with the Pt particle layer in order to establish contact with the underlying Au NW tracks. After aligning the Au NW tracks with the electrode layer on an x, y, z stage, they were brought in contact and a force was applied to press the Pt particle electrodes through the top semi-cured PDMS layer and into contact with the Au NW tracks. The force must be tuned in order to achieve both good contact between the electrodes and tracks and the desired height for the final electrodes. The device was then cured for 20 minutes at 80°C before removing the electrode layer membrane by dissolving the photoresist with acetone and fully curing the device overnight at 80°C. The implant was cut in shape with a scalpel and carefully removed from the wafer. It was then mounted on a custom PCB by stencil printing silver epoxy on the contacts and securing it with epoxy glue. An Omnetics nano connector (part number A76284-001) was soldered to the other side of the PCB to minimize the size and weight of the device.

#### Device characterization

All electrochemical measurements prior to implantation were conducted with a potentiostat (Autolab, Metrohm), in a 3-electrode configuration with an Ag/AgCl reference electrode, a Pt wire counter electrode and PBS as electrolyte. Before every impedance measurement, six voltammetry cycles between −0.6 – 0.8 V were run at 0.2 V/s on each electrode to assure a stable interface. The impedance measurements were taken in the range of 1-100 kHz at 50 points. The equivalent circuit was determined and fitted with the Nova 2.1.2 software.

#### Preparation for implantation

Prior to implantation (within 30 minutes), each array was conditioned with a procedure to improve *in vivo* performance and increase signal by establishing a stable electrode double layer and vacate air trapped in the porous electrode structure. The conditioning protocol consisted of voltage sweeps between −0.6 – 0.8V for at least 20 repetitions.

#### Animals and surgical procedures

All methods in mice were carried out according to the guidelines of the Veterinary Office of Switzerland and following approval by the Cantonal Veterinary Office in Zurich (license 133/2015). A total of 8 adult male mice (4-10 months old) were used in this study. Mice were triple transgenic SNAP25-Cre;CamK2a-tTA; TITL-GCaMP6f animals, expressing GCaMP6f in excitatory neurons. To generate triple transgenic animals, double transgenic mice carrying CamK2a-Tta^[35]^ and TITL-GCaMP6f^[31]^ were crossed with a SNAP25-Cre line^[36]^ individual lines are available from The Jackson Laboratory as JAX# 016198, JAX#024103, and JAX# 022864, respectively). Mice were anesthetized with isoflurane (in pure O_2_, 2% for induction, 1.5% for surgery) and body temperature was maintained at 37°C. Local analgesia (lidocaine 1%) was applied topically, and the skull was exposed and cleaned to provide access the entire dorsal surface. A thin layer of adhesive (iBond; UV-cured) was distributed across the dry surface of the skull and dental acrylic (Charisma) was used to form a wall around the left hemisphere. A thin layer of dental cement was then distributed homogenously over the exposed skull (Tetric EvoFlow A1). Next, a small hole (< 0.5 mm) was drilled over the cerebellum to allow placement of the reference electrode. We performed a 5-mm diameter craniotomy, centered over the left somatosensory cortex. The dura mater was removed while irrigating with Ringer solution to avoid drying or heating. The ECoG was held with a stereotaxic arm and positioned over the craniotomy. Once the ECoG was positioned, the connector PCB was secured to the edge of the skull, atop the acrylic wall with additional dental cement. The ECoG array was then held in place by a glass capillary covered by a blunt, plastic pipette tip. While the ECoG was held flush with the surface of the brain with gentle pressure, dental cement was applied to hold it in position flush with the skull. A small PDMS well was secured to the ECoG with additional PDMS layer applied at the interface to allow imaging through the ECoG with a water-immersion objective. The reference was placed in the hole over the cerebellum and fixed with dental cement. Finally, a metal post for head fixation was affixed to on the back of the right hemisphere with dental cement. Mice were housed individually after implantation to avoid damage to each other or the implant.

#### ECoG recording from Opto-e-Dura

Mice were head-fixed after habituation to restraint. The array was connected to a multiplexing headstage and multichannel amplifier (RHD2000 Intan) and *in vivo* impedance was measured, after which ECoG signals were recorded onto a desktop PC. ECoG signals were sampled at 20 kHz and offline filtering was performed. ECoG data were low pass filtered with a second order Butterworth filter below 200 Hz and down sampled to 1 kHz. For recordings under anesthesia, animals were first anesthetized using isoflurane (1%) and body temperature was maintained at 37°C using a heating pad.

#### Wide-field calcium imaging

To demonstrate the suitability of the Opto-e-Dura for combination with optical methods, we first performed wide-field calcium imaging from the dorsal cortex while simultaneously acquiring electrophysiological signals from the ECoG. The wide-field imaging system and methods have been described previously^[28]^. Briefly, a sensitive CMOS camera (Hamamatsu Orca Flash 4.0) was mounted on top of a dual objective setup. Two objectives (Navitar; top objective: D-5095, 50-mm focal length, f/# 0.95; bottom objective inverted: D-2595, 25-mm focal length, f/# 0.95) were interfaced with a dichroic mirror (510 nm; AHF; Beamsplitter T510LPXRXT) inside a filter cube (Thorlabs). This combination allowed a ∼9 mm field-of-view, covering most of the dorsal cortex of the one hemisphere. Blue LED light (470 nm; M470L3; Thorlabs) was guided through an excitation filter (514/30 nm BrightLine HC) and a diffuser, collimated, reflected from the dichroic mirror, and focused through the bottom objective approximately 100 µm below the blood vessels. Green light emitted from the preparation passed through both objectives and an emission filter (515/30 nm) before reaching the camera. The total power of blue light on the preparation was <5 mW, i.e., <0.1 mW/mm^2^. At this illumination power we did not observe electrical artifacts on the ECoG, nor photobleaching of the fluorescent indicator. Data was collected with a temporal resolution of 20 Hz and a spatial resolution of 512×512 or 2048×2048 pixels. On each imaging day a green reflectance image was taken (fiber-coupled LED illuminated from the side; Thorlabs) as reference to capture blood vessel patterns.

#### Functional localizer for mapping of the cortex covered by the Opto-e-Dura

Each mouse underwent a mapping session under anesthesia (1% isoflurane), in which we presented four different sensory stimuli: tactile stimulation with a moving bar touching either multiple whiskers, or the forelimb or the hindlimb (25 Hz for 2 s), and visual stimulation with a blue LED in front of the eye (100-ms duration flashes). The averaged evoked maps showed activation patches in the expected areas^[33]^ (**Figure 4**c), which were used to define the general borders of sensory areas. These locations allowed us to confirm the location of the array on the brain, and provided a basis for assessing the correspondence between electrical signals recorded on the ECoG array and in the bulk fluorescence signals of the calcium indicator.

#### Two-photon calcium imaging

In two mice, we acquired two-photon imaging data. We used a Scientifica two-photon microscope controlled by ScanImage, equipped with a Ti:sapphire laser (Mai Tai HP; Newport Spectra Physics), a water-immersion objective (Nikon 16x LWD-PF, NA 0.8,3-mm working distance), galvanometric scan mirrors, and a Pockel’s Cell (Conoptics) for laser intensity modulation. We excited GCaMP6f at 940 nm and detected green fluorescence with a photomultiplier tube (Hamamatsu). Images (256×128 pixel) were acquired at 3-Hz frame rate and tens of cells per field of view were imaged simultaneously.

#### Linear electrode recordings

To test penetration of the Opto-e-Dura with a sharp linear electrode array, we used a 32-contact silicon array (15 micron Iridium Oxide contacts and 50 micron site spacing, model E32-50-S1-L6 NT, Atlas Neurotechnologies). Prior to insertion, the array was dipped in a lipophilic fluorescent dye (DiD molecular probes) to enable post-mortem localization. The array was subsequently attached to a mechanical micro-drive (Narishige) and inserted through the ECoG and into the brain. Data from the 32-contact linear array were recorded by a second headstage connected to the same multichannel amplifier used to record the ECoG (RHD2000 Intan). Recordings were made at various locations through the Opto-e-Dura, both adjacent to ECoG electrodes and in the center of the array. After awake recordings were performed, additional measurements were made in anesthetized animals with isoflurane (1%). During dual recordings with ECoG and linear arrays, all electrophysiological data were acquired at 30 kHz and filtered into low and high frequency activity offline. Both ECoG and LFP data were low pass filtered with a fourth order Butterworth filter below 200 Hz and down sampled to 1 kHz. The data from the linear array were additionally filtered into a high frequency component using a fourth order Butterworth bandpass (640 – 6000 Hz). Single unit activity was extracted by a taking only the largest unit with amplitude at least larger than 5 times the standard deviation of the high pass signal.

#### Data analysis

Data analysis was performed using Matlab software (Mathworks).

Wide-field fluorescence images were sampled down to 256×256 pixels and pixels outside the imaging area were discarded. Each pixel and each trial were normalized to baseline several frames before the first auditory cue (frame 0 division). To study neural dynamics during sensation, we used a different baseline, several frames before the relevant stimulus. Electrical and optical signals were processed using in-house software. For comparison of ECoG and calcium signals, ECoG was decomposed into time-resolved power envelopes using the fast Fourier transform with a Hann-tapered window with a length dependent on the frequency (5 cycles) and a temporal resolution to match the simultaneous calcium recording. Calcium time-series and the band-limited enveloped from the ECoG were then correlated using the Spearmann rank correlation because both power and ΔF/F are strictly positive. Spike locking was computed between single units isolated as described above and the simultaneous ECoG signal using the pairwise phase consistency metric^[37]^. The Fourier components of ECoG data centered on spikes were computed in frequency-dependent windows (5 cycles) after applying a Hanning taper. The mean length of the resultant vector from all pairs of Fourier components for a given unit provide a metric of the phase locking.

## Acknowledgements

We would like to thank Dr. Ladan Egolf for transgenic mice, Drs. Rolf Pfister and Peter Hamm for spectrometry, ScopeM for providing access to the SEM and Martin Lanz, Aldo Rossi and Stephen Wheeler for technical support, Lazar Shumanovski for help with histology, and the members of the Helmchen Lab for their help and advice during the project. We acknowledge financial support by ETH Zurich (J.V.), and the Swedish Government Strategic Research Area in Materials Science on Advanced Functional Materials at Linköping University (Faculty Grant SFO-Mat-LiU No. 2009-00971, K.T.), the Swedish Foundation for Strategic Research (K.T.), a Forschungskredit from the University of Zurich (project K-41220-04, C.L.), and the European Research Council (ERC Advanced Grant BRAINCOMPATH, project 670757; F.H.)

## Supporting Information

**Figure S1.**
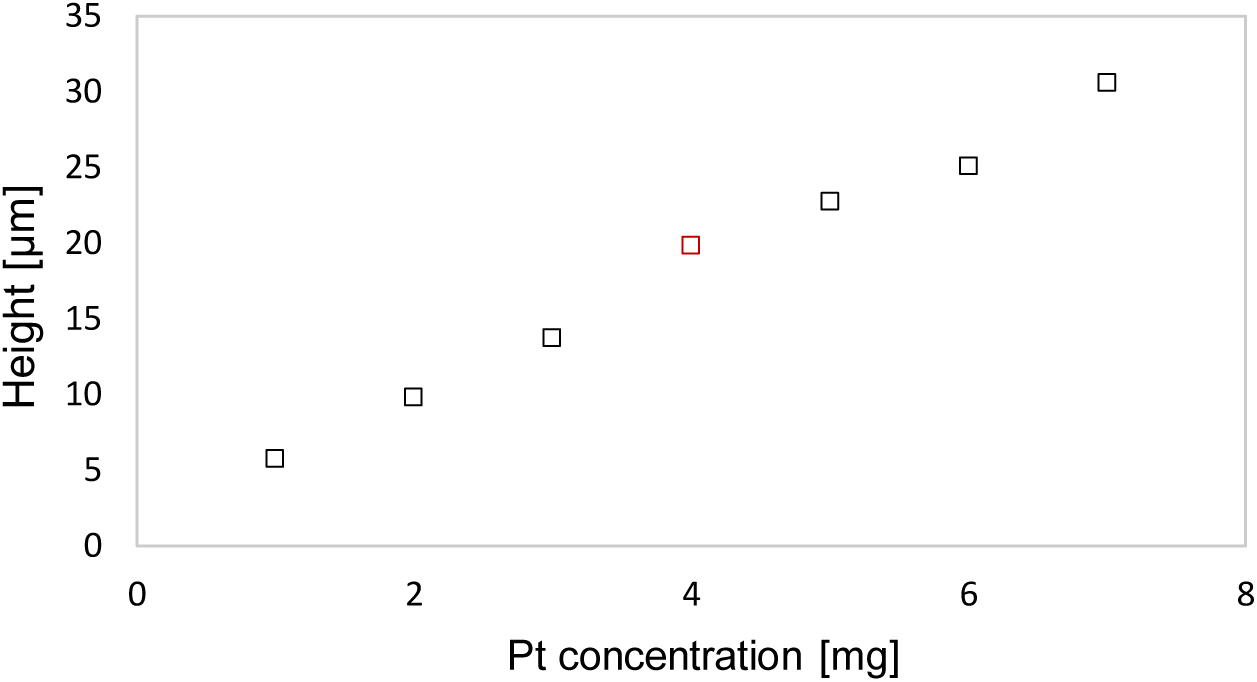
Determination of Pt pillar height regarding the various Pt particle concentrations. For the here presented electrodes a concentration of 4 mg per surface area was chosen to gain electrode pillars of roughly 20 µm.

**Figure S2.**
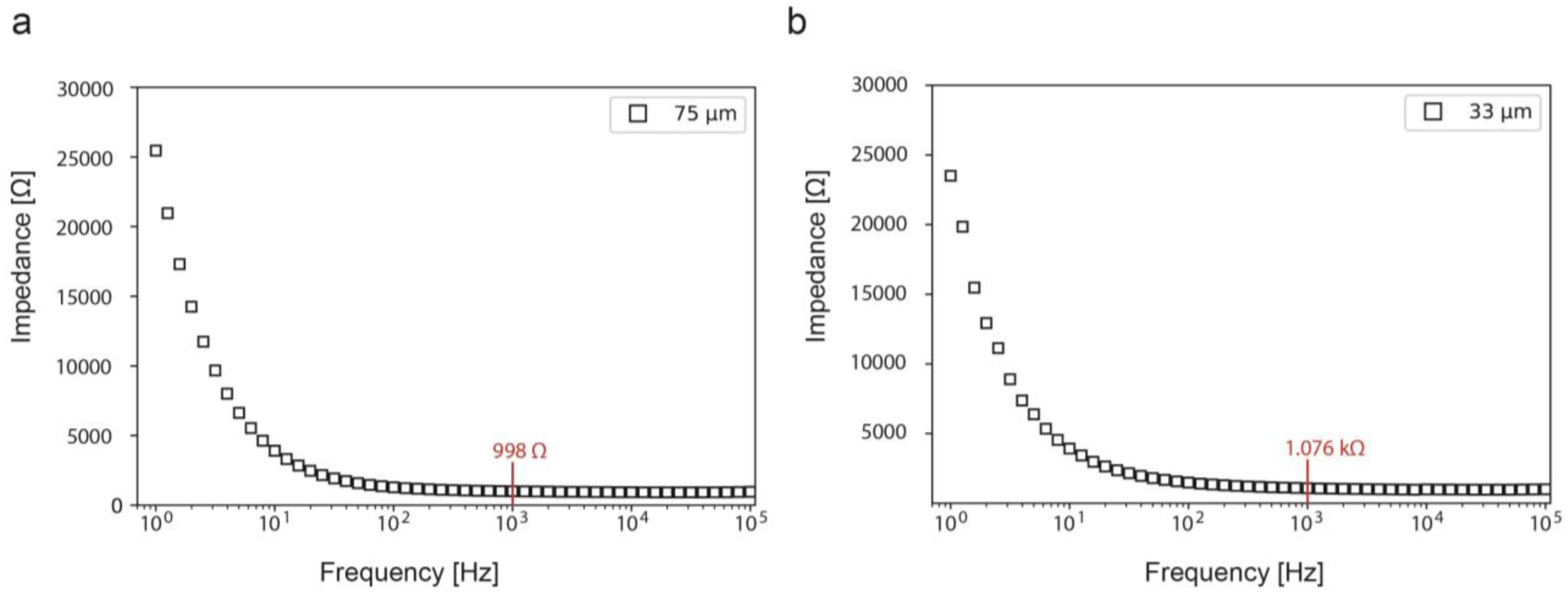
Impact of full ECoG device thickness on the impedance a) 75 μm thick and b) 33 μm thick ECoG arrays. The data shows the median and the upper and lower quartile.

**Figure S3.**
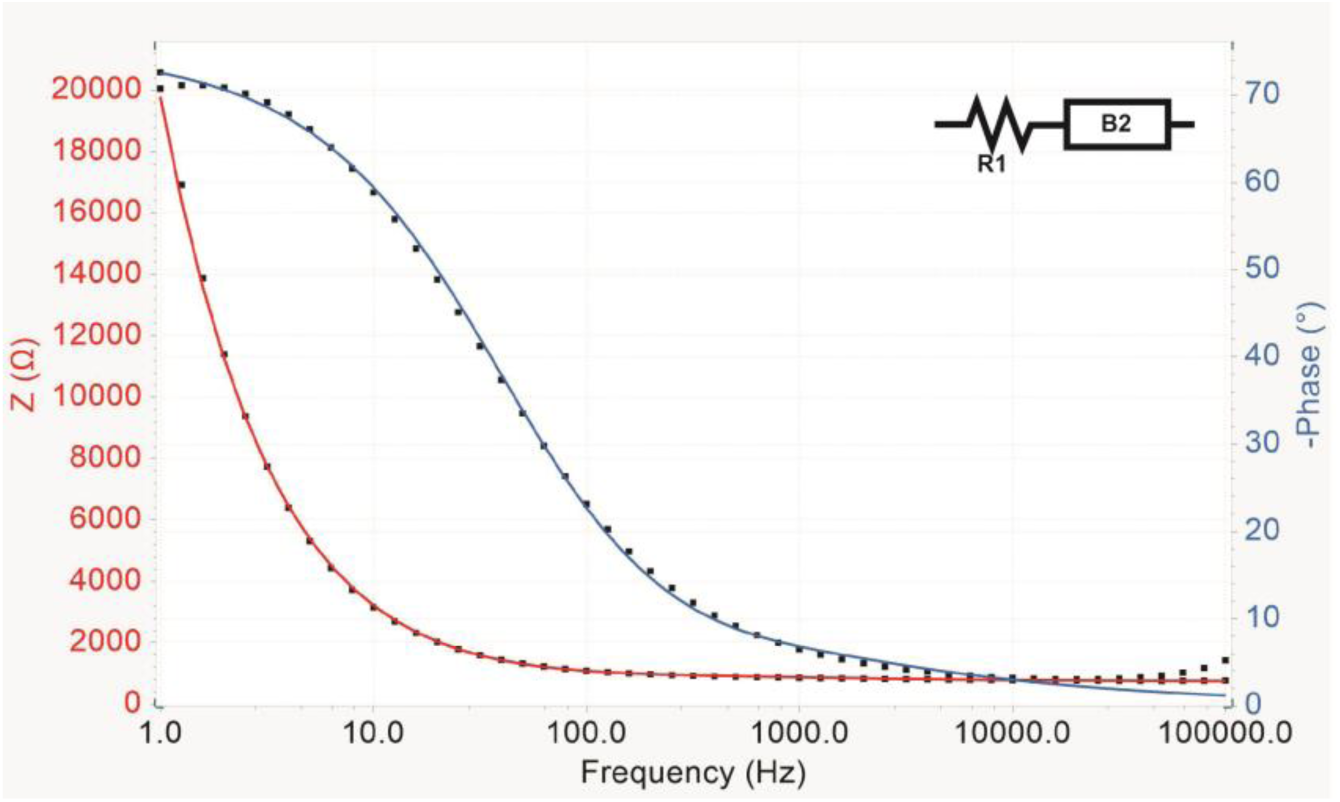
Equivalent circuit for the ECoG electrodes fitted in blue for the phase and red for the impedance to the corresponding black measurement values. The circuit consists of R1 (730 Ω), the resistance for electronics and electrolyte in series with an transmission line model from Bisquert^[38]^ with R2 (24.8 MΩ) the electrolyte resistance of the pore, R3 (740 MΩ) accounting for the pore resistance and Q (553 mMho*s^0.834^) the constant phase element, while L, indicating the pore length was set to 20E-6. However, due to the complexity of the porous electrode-electrolyte it is a simplified circuit.

**Figure S4.**
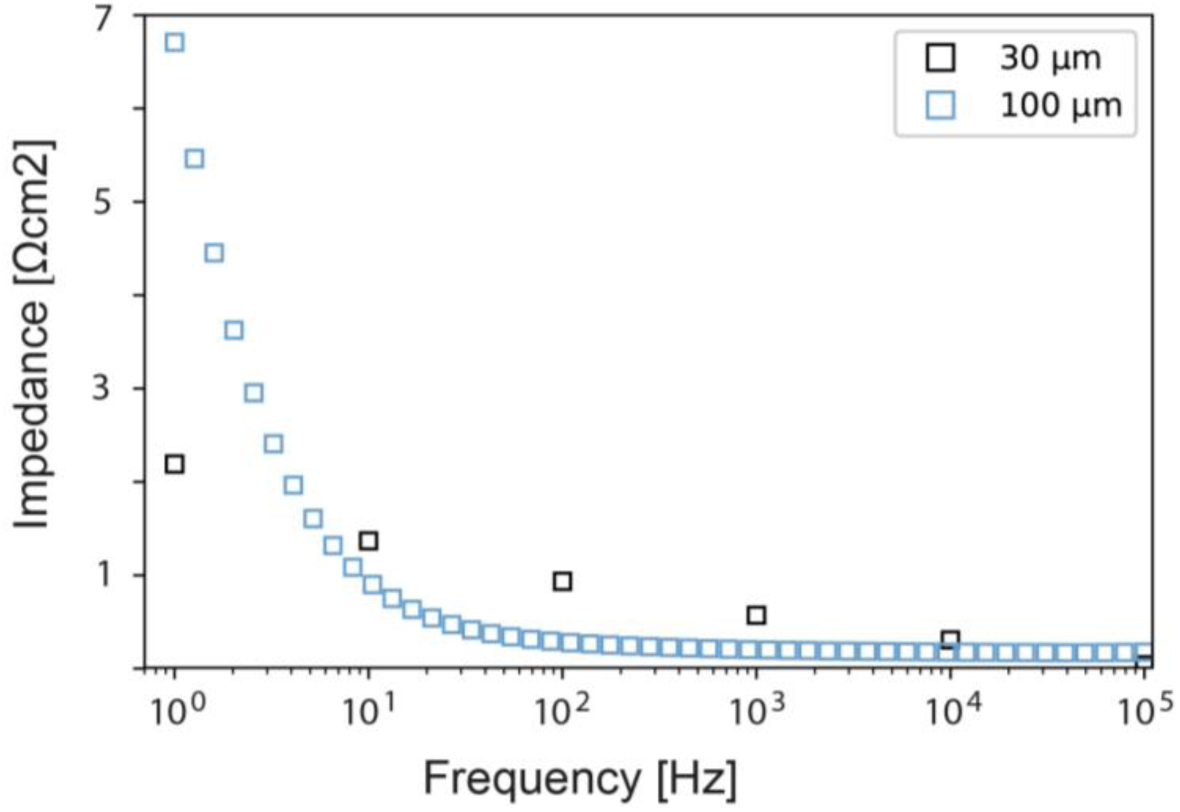
Due to the high surface area of porous Pt electrodes a significantly decreased impedance to area performance was achieved with the here presented fabrication technique. This was the case for electrode of A) 100 μm diameter with roughly 6.66 Ωcm2 at 1 Hz and B) electrodes with roughly 30 μm in diameter achieving 1.14 Ωcm2 at 1Hz, being the lowest value documented for stretchable electrodes.

**Figure S5.**
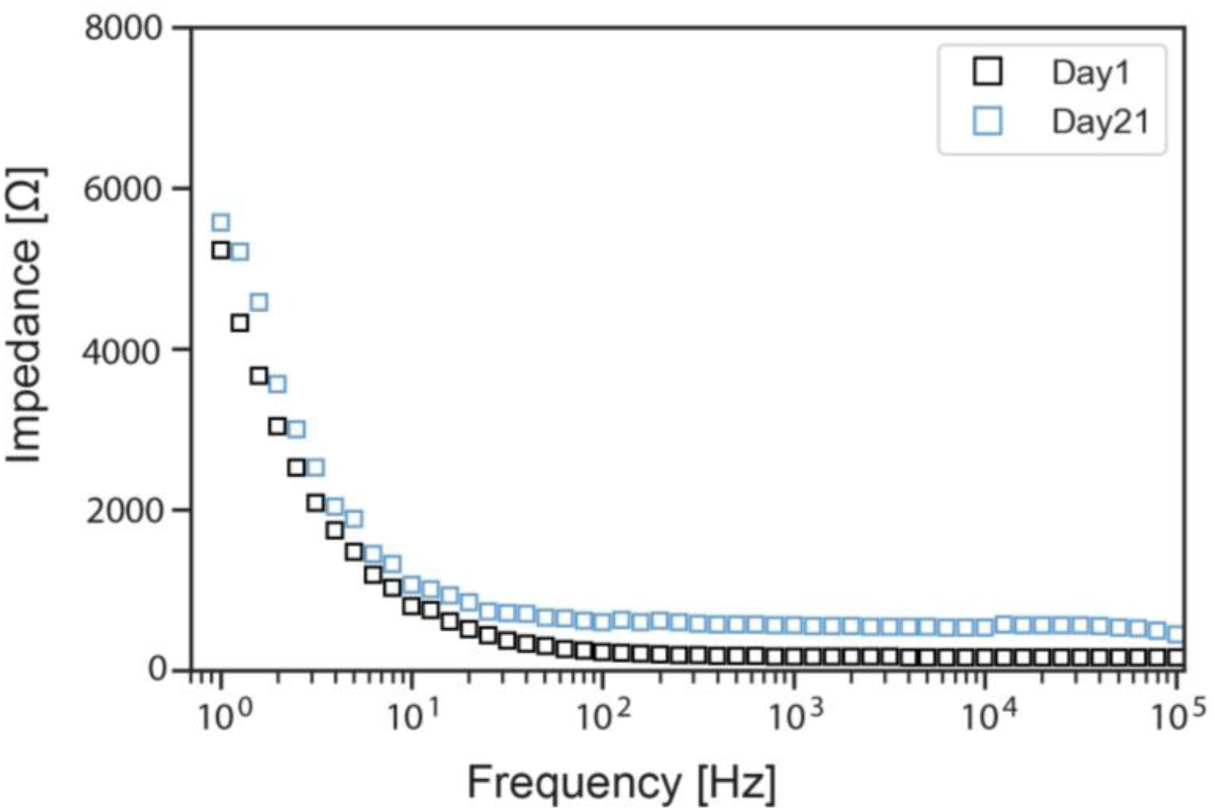
Stability test of electrodes for 21 days at 85°C in PBS, with impedance data from day 1 (black) and day 21 (blue). The increased impedance in day 21 can be explained by an overall higher electrolyte impedance, as it is a constant increase over all frequencies.

**Figure S6.**
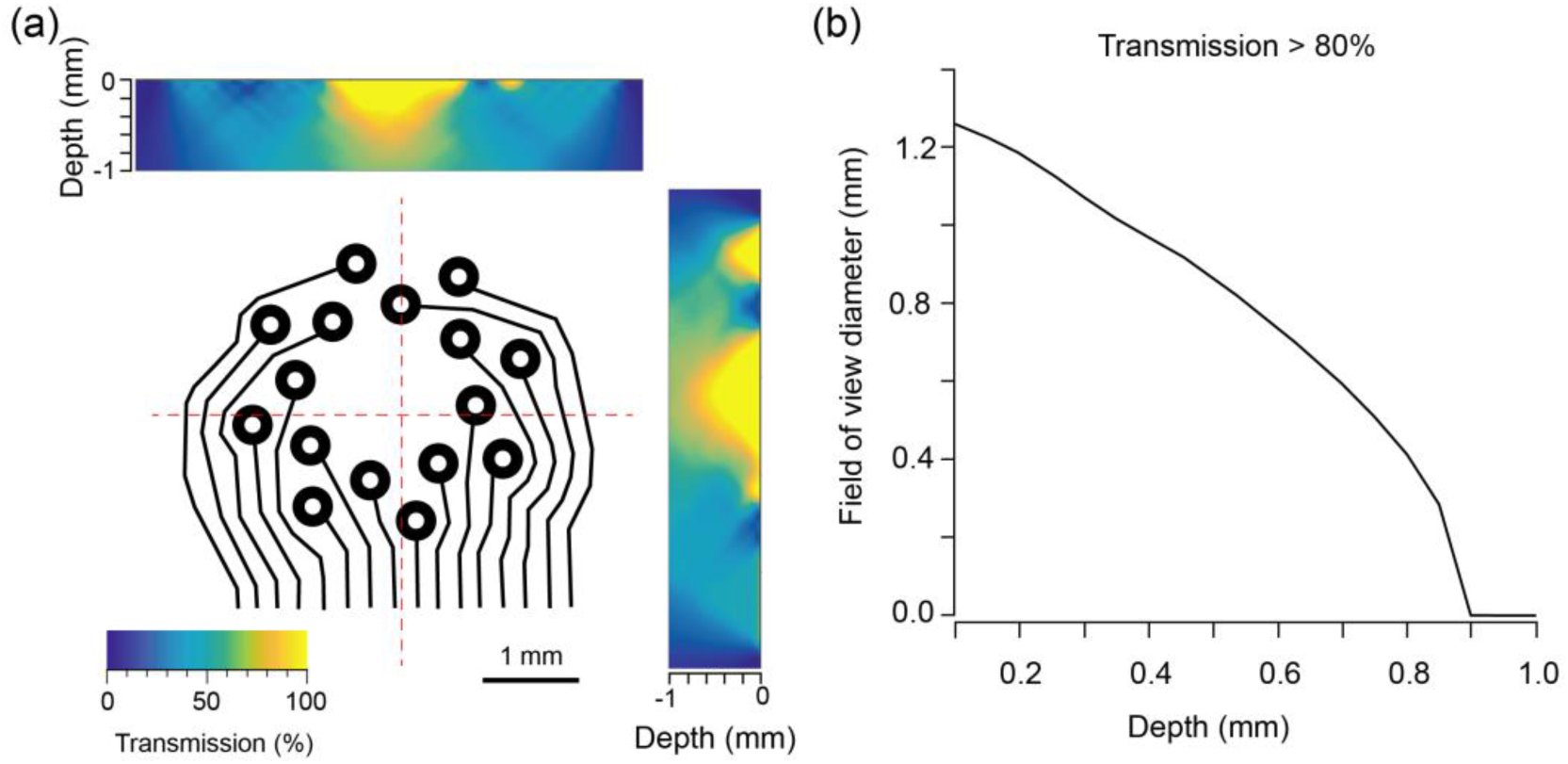
Estimation of signal loss and field of view when imaging through the opto-e-dura. a) Estimated transmission of light through two cross sections of the opto-e-dura. The array is depicted in the center panel, with cross-sections indicated in red. Top, estimate of light transmission in a horizontal cross section of the array as a function of focal depth relative to the array. Right, estimate of light transmission in a vertical cross section of the array as a function of focal depth relative to the array. b) Estimated field of view diameter with transmission greater than 80% at depths from 0.1 to 1 mm.

